# A mathematical approach for secondary structure analysis can provide an eyehole to the RNA world

**DOI:** 10.1101/079608

**Authors:** Nikolaos Konstantinides

**Affiliations:** Department of Mathematics, University of Crete, 71409 Heraklion, Crete, Greece; Current address: Department of Biology, New York University, 1009 Silver Center, New York, NY, USA

**Keywords:** pseudoknot, secondary structure, analytic combinatorics, neutral network, sequence to structure map, molecular evolution, robustness, evolvability

## Abstract

The RNA pseudoknot is a conserved secondary structure encountered in a number of ribozymes, which assume a central role in the RNA world hypothesis. However, RNA folding algorithms could not predict pseudoknots until recently. Analytic combinatorics – a newly arisen mathematical field – has introduced a way of enumerating different RNA configurations and quantifying RNA pseudoknot structure robustness and evolvability, two features that drive their molecular evolution. I will present a mathematician’s viewpoint of RNA secondary structures, and explain how analytic combinatorics applied on RNA sequence to structure maps can represent a valuable tool for understanding RNA secondary structure evolution. Analytic combinatorics can be implemented for the optimization of RNA secondary structure prediction algorithms, the derivation of molecular evolution mathematical models, as well as in a number of biotechnological applications, such as biosensors, riboswitches etc. Moreover, it showcases how the integration of biology and mathematics can provide a different viewpoint into the RNA world.

## Introduction

The RNA world hypothesis states that RNA is the first molecule to ever be able to store genetic information itself and, in the same time, have regulatory and catalytic activities ^1^. DNA and proteins later assumed the tasks of carrying the genetic information and catalyzing chemical reactions, respectively. These RNA molecules, that act as enzymes and are also capable of carrying genetic information to replicate themselves, are called ribozymes. Since their discovery ^2^, many ribozymes have been reported ^3–5^, and one common feature is their complex secondary structure ^6^. Understanding how ribozymes implement their double nature can help us explain how the RNA-based world evolved to then give rise to the DNA and protein-centered world that we live in.

RNA molecules assume various structures - some of which rather different from the minimum free energy arrangement ^7^- depending on relatively unknown conditions. The catalytic activities of any RNA molecule depend on it structure ^8^; hence, different structures of the same sequence perform diverse functions ^9^. Moreover, the structure of an RNA molecule provides information about its genetic robustness and evolvability ^10^. A functional structure has to achieve a balance between its capacity to withstand and its flexibility to allow change when faced with structure-altering mutations. This can be achieved if the same structure may arise by different sequences. Depending on how these different sequences are distributed in the sequence space, a structure can be more or less robust and evolvable ^11^. Evidently, RNA structure, function and evolution are tightly correlated.

Analytic combinatorics – a newly arisen mathematical tool - has been used lately to study RNA structures, as exemplified here in the case of RNA pseudoknots. I present here how the combination of analytic combinatorics with RNA sequence to structure maps and their neutral networks can provide new insight about the evolutionary history of different RNA structures offering, at the same time, new parameters towards a more complete mathematical model of evolution ^12^.

## The RNA pseudoknot is a conserved conformation of specific RNA molecules

The most common RNA structures consist of non-mutually crossing base pairs; these are defined as hairpin structures (Fig. 1A). RNA pseudoknots are formed when ribonucleotides that belong to the loop of the hairpin base-pair with ribonucleotides that do not belong to the hairpin (Fig. 1A). An RNA structure that contains one or more pseudoknots is defined as a pseudoknot structure. Furthermore, pseudoknot structures can be divided in ordinary and non-ordinary ones. Non-ordinary pseudoknots are more complex structures, rarely observed in nature (http://pseudobaseplusplus.utep.edu) and will only be briefly discussed (Fig. 1A). Obviously, both hairpin and pseudoknot structures can vary a lot in terms of ribonucleotide number, hairpin conformation, base-pairing topology etc, all these affecting their structure and function.

**Figure 1 legend:**
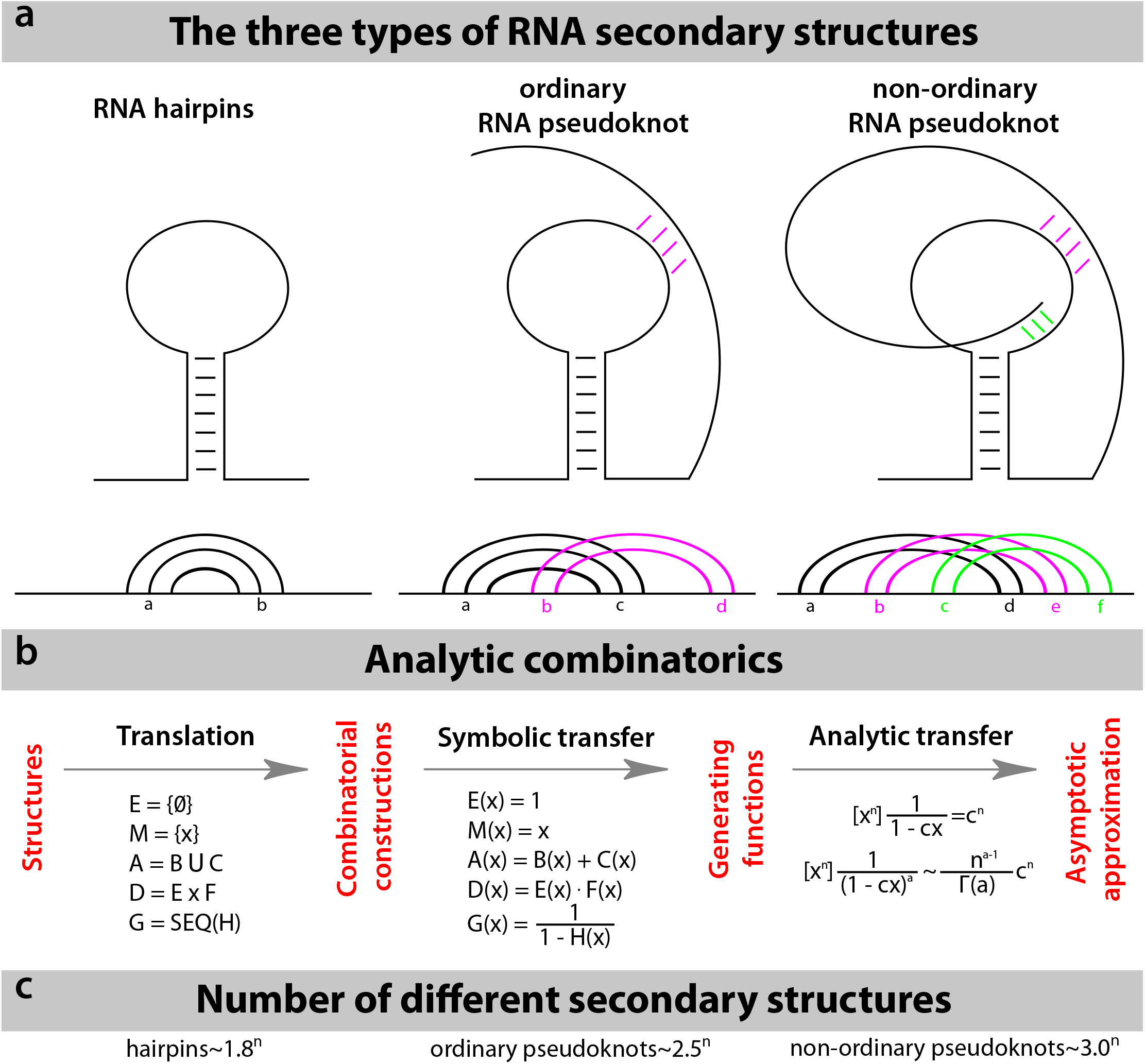
Enumeration of structures with analytic combinatorics. (A) In terms of base-pairing, a string of ribonucleotides can acquire three different forms. The most common one is the hairpin, where no mutually crossing base-pairs are observed. Underneath the structure, the RNA molecule is illustrated as a straight line, whereas curved lines represent base pairs between specific ribonucleotides, highlighting the absence of mutually crossing base pairs. The ordinary RNA pseudoknot is a structure that forms when a single-stranded nucleotide stretch of a hairpin loop base-pairs with nucleotides elsewhere in the same molecule. Nucleotide string *a* base-pairs with nucleotide string *c* and nucleotide string *b* with nucleotide string *d,* where *a < b < c < d.* Non-ordinary, more complex pseudoknot structures are rarely observed in nature. These can carry more than two mutually crossing base pairs. (B) Using analytic combinatorics, one can approximate the number of different structures. Initially, the structures are translated into combinatorial constructions, using natural combinatorial operators, such as union (set A), Cartesian product (set D) and sequences (set G), between operands, such as empty sets (set E), unit sets (set M) or other constructions. Subsequently, through symbolic transfer equations, generating functions can be formed. The *n*^th^ coefficient of a generating function *g*(*x*) (represented as [*x*^*n*^]*g*(*x*)) can, in turn, be approximated through the use of analytic transfer equations. (C) The number of different structures of each of the aforementioned types of RNA secondary structure has been approximated using analytic combinatorics.

Several molecules acquire ordinary pseudoknot structures ^13, 14^. Most of their structures and functions are conserved amongst all the organisms in which they are found, although their primary sequence may vary. It is rather intriguing to observe that most, if not all, RNA components of functional ribozymes exhibit a pseudoknot structure (23S rRNA, RNase P, Telomerase, group I introns, hepatitis delta virus ribozyme etc). Ribozymes assume an important role in the RNA world hypothesis, stating that biological systems have their origins in self-replicating molecules made of RNA; therefore, the evolution of these ribonucleoprotein complexes has received much attention ^15, 16^. Only recently, the prediction of pseudoknot structures by RNA structure prediction algorithms has become possible ^17^. Lately, mathematical biologists have studied pseudoknots extensively using analytic combinatorics ^18–20^. Using their calculations, which are presented here, we may obtain information about the evolution of RNA secondary structures.

## Neutral networks of RNA structures

To achieve this, we need to study the neutral networks of different RNA structures. Sequence to structure maps are valuable for studying RNA evolution. These maps consist of two sets, the set of sequences and the set of structures. Each sequence corresponds to one structure. Since sequences are more than the structures, many members of the sequence set are connected with one member of the structure set. The sequences that are connected to one structure form a subset, which is called neutral set. Neutral networks are the neutral sets in which all members are connected with each other with successive single mutations that do not lead to sequences with different structure (neutral neighbors). In the case of RNA sequence to structure maps, neutral sets are always coherent and, therefore, the two terms are interchangeable ^21^.

The magnitude and the shape of the neutral network of a given structure can be correlated with its robustness - the capacity of the structure to resist change retaining its configuration - and its evolvability - its liability to change. Although, in nature, the sequences of a neutral network are not equally distributed, it is safe to assume that the larger a neutral network the more robust the structure, since single mutations are more likely to result in retention of the original structure.

It is intuitive to perceive robustness and evolvability as two opposite traits, since they describe the structure’s resistance and liability to change, respectively. However, structure robustness is positively correlated to structure evolvability ^10^; a structure that is robust has a more extensive neutral network, which, in turn, neighbors more structures, enhancing the evolvability of the structure. System drift on a large genotype network is necessary for the production of novel phenotypes, as this drift despite keeping the phenotype intact alters the neighborhood of the structure, potentially bringing it one mutation away from a novel structure ^10, 22, 23^. Therefore, a larger network is not only robust but also more evolvable by increasing the amount of different accessible phenotypes. The capability to evaluate the magnitude and/or the shape of the neutral network of a structure can offer an additional criterion when assessing its evolutionary history.

## Analytic combinatorics: another tool in our mathematical toolbox

In this article, an RNA structure is defined as a molecule with a specific topology of base pairs between its ribonucleotides. If the identity of the ribonucleotides, whether forming base pairs or not, changes, a different (new) structure will not be considered to have formed as long as the base-pairing topology remains the same.

### Theory

Analytic combinatorics ^24^ is a clade of mathematics used to estimate specific properties of large configurations, as - in the present case - the number of different structures an RNA molecule of a given size can acquire. The main mathematical tool is the generating function, which is described next. Analytic combinatorics unites two different disciplines. First, it uses combinatorics to translate the examined configuration into a generating function. Subsequently, analytical methods are deployed to extract information from these functions.

The functions we are familiar with are of the type *y* = *f*(*n*), where *y* can be easily determined by substituting *n* with the desired value. For instance, the formula for calculating the different sequences of RNA molecules of length *n* is the following:

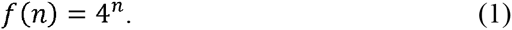

However, for most biological systems a simple explicit formula is impossible to be computed; therefore, to describe such systems, generating functions can be a valuable alternative. Generating functions are power series

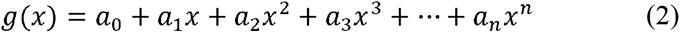

whose coefficients *a*_*n*_ are the answers to the problem in hand. This means that the *a*_*n*_ coefficient of a generating function corresponds to the *f*(*n*) of an explicit formula.

Generating functions should be used to approach a problem only if no explicit formula is available; they are not easy to handle, since the evaluation of the n^th^ coefficient of the function (when *n* is large) can be very demanding and often can only be approximated.

### Process

An important part of this analysis is the translation of the configuration to be studied to a combinatorial construction. This is performed using simple operators such as unions, Cartesian products and sequences, and operands such as unit sets (sets with only one member) or other constructions (Fig. 1B), as exemplified subsequently.

Afterwards, the generating function can be calculated directly from the combinatorial construction, through an algorithm that translates the operands and the operators into mathematical expressions. The theorems, upon which these translations are founded, are called symbolic transfer theorems (Fig. 1B).

Finally, the transfer from the generating function to the estimation of its coefficients relies upon a third algorithm. Depending on the type of the generating function, this algorithm utilizes certain theorems (analytic transfer theorems) to achieve the approximation of the coefficients of the generating function (Fig. 1B). The transfer theorems, the analytic ones in particular, are difficult to prove but easy to implement. Many transfer theorems are already available ^24, 25^ and the proof of new ones is an active area of study in analytic combinatorics.

### Example

One can calculate the generating function of the different sequences of RNA molecules of length *n*, based on the process described earlier and the theorems summarized in Fig. 1B, and compare the result with the already known explicit formula (1).

The set of the ribonucleotides R (a 4-member set consisting of the four possible ribonucleotides) is the union of the four unit sets, one of each nucleotide. Hence, its generating function is equal to four times the generating function of the unit set (*M*(*x*) = *x*):

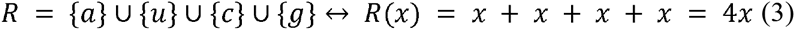

The members of the set of the different sequences of RNA molecules, S, are sequences of members of the ribonucleotide set, therefore:

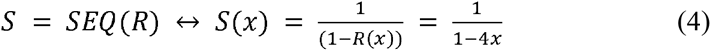

It is, afterwards, easy to verify that:

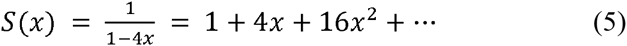

and compare the results of the explicit function (1) with the coefficients of the generating function (5):

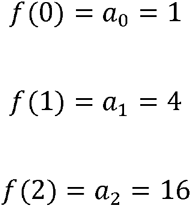

etc.

The calculated generating function (4) is of the type:

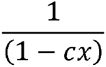

where *c* = 4. Therefore, the corresponding analytic transfer theorem as indicated in Fig. 1B is the following:

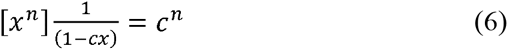

where 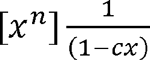 symbolizes the *n*^th^ coefficient of the generating function 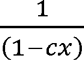. We can deduce that the coefficients of *S*(*x*) can be approximated by the following function:

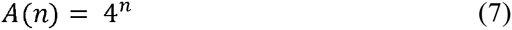

which, in the present example, equals the exact number of the different sequences of RNA molecules of length *n.*

The most intriguing part of analytic combinatorics is that it creates an interface of collaboration between different disciplines - mathematicians and biologists – and can be used to exploit beautiful mathematics, such as analysis, to assess exciting evolutionary questions. Nevertheless, it can also be helpful for biologists without mathematical background who can use the algorithm by simply applying proven transfer theorems.

## Neutral networks and evolution of RNA pseudoknots

The use of analytic combinatorics for the enumeration of RNA configurations was introduced in the early ‘80s ^26^. Recently, the number of different structures of hairpins and pseudoknots was approximated ^27–29^ (Fig. 1C), indicating that ordinary pseudoknot structures exhibit large neutral networks. Mutational robustness of large neutral networks offers an apparent selective advantage ^21^. Moreover, the base-pair number in pseudoknot structures follows a Gaussian distribution ^30^, designating that the majority of pseudoknot structures have a similar number of base pairs.

The magnitude of neutral networks is inversely proportional to the number of different structures. Having this in mind as well as the numbers of different structures of each RNA type (Fig. 1C), one can estimate the extent of neutral networks and, hence, the robustness and evolvability of each different structure. Hairpins are, on average, more robust than ordinary pseudoknots, which in turn are more robust than the non-ordinary pseudoknots. Inversely, the ordinary pseudoknots can acquire more different structures than the hairpins. Analytic combinatorics could generate information about the relationship between robustness and evolvability of structures (robustness is influenced by the magnitude of the neutral networks, whereas evolvability depends on both magnitude and shape). Combined with research about the distribution of sequences that fold into a given structure ^17, 31, 32^, these studies can enhance our understanding regarding the production of evolutionary innovations in a robust environment.

The mathematical framework presented earlier allows us to survey the prevalence of ordinary pseudoknot structures in ribozymes from a different viewpoint. Robustness and evolvability of RNA pseudoknots, alongside their presence in every known ribozyme, concur with the RNA world hypothesis, which requires robust RNA molecules able to acquire diverse structures. These neutral network characteristics are ideal for the rapid evolution of robust structures.

## Future applications for RNA structure and molecular evolution studies

Being able to assess the number of different structures of a given set of RNA molecules can assist the development of RNA secondary structure prediction algorithms. In this context, the set of biophysically relevant pseudoknot RNA structures (which satisfy certain minimum free energy prerequisites) has been enumerated, indicating this small set as a potential output class for future prediction algorithms ^27^. Moreover, consideration of the magnitude of a structure’s neutral network, as a meta-prediction procedure in RNA secondary structure computation algorithms, could serve as validation of the output structure. The aforementioned approach could thus generate one of the missing pieces for comprehending the principles of RNA folding. Moreover, since the implementation of algorithms for the prediction of the joint structure of two interacting RNA molecules, combinatorial analysis of interacting RNA molecules has been performed ^33^.

Mutations occur in the level of genotype (sequence), while selection acts on the phenotypic output (structure). Therefore, the derivation of mathematical models that describe evolution as accurately as possible requires the integration of accurate representations of genotype-phenotype maps. To add on this complexity, the phenotype may be manifested in different levels, from molecules to organelles, cells, tissues, organs, and organisms. Analytic combinatorics offers an alternative way of exploring the fitness landscape by evaluating the robustness and evolvability of different phenotypes.

Lately, a number of biotechnological applications have resided in the use of ribozymes as biosensors for environmental monitoring ^34, 35^, as riboswitches for genetic engineering etc ^36–38^. Many synthetic ribozymes have been produced using experimental evolution ^39^. Having a way to evaluate the robustness and evolvability of ribozyme structures is critical for the rational design of RNA molecules. Finally, understanding how ribozymes have evolved to exert their functions, we will be able to engineer catalytic RNA molecules, whose function may be controlled using chemical or physical signals, more efficiently ^37, 40^.

## Conclusion

Analytic combinatorics has offered a plausible explanation as to why RNA pseudoknot structures were recruited and retained until today in fundamental biological molecules and procedures. Analytic combinatorics can provide the foundation, upon which biologists can approach previously inaccessible fields of research. Moreover, by describing the landscape of RNA sequence to structure maps analytic combinatorics presents the opportunity to cover some of the gaps towards a mathematical model of evolution ^12^.

## Acknowledgements

I thank M. Kolountzakis for introducing me to Analytic Combinatorics and M. Averof and A. Papantonis for comments on the manuscript.

